# Near Zero Photon Bioimaging

**DOI:** 10.1101/2024.06.12.598699

**Authors:** Lucas Sheneman, Sulaimon Balogun, Jill L. Johnson, Maria J. Harrison, Andreas E. Vasdekis

## Abstract

Enhancing the reliability and reproducibility of optical microscopy by reducing specimen irradiance continues to be an important biotechnology target. As irradiance levels are reduced, however, the particle nature of light is heightened, giving rise to Poisson noise, or photon sparsity that restricts only a few (0.5%) image pixels to comprise a photon. Photon-sparsity can be addressed by collecting more than 200 photons per pixel; this, however, requires extended acquisition durations and, thus, suboptimal imaging rates. Here, we introduce near-zero photon imaging, a method that operates at kHz rates and 10,000-fold lower irradiance than modern microscopy. To achieve this performance, we deployed a judiciously designed epi-fluorescence microscope enabling ultralow background and artificial intelligence that learns to reconstruct biological images from as low as 0.01 photons per pixel. We demonstrate that near-zero photon imaging captures the structure of both multicellular and subcellular targets with high fidelity, including features represented by nearly zero photons. Beyond optical microscopy, the near-zero photon imaging paradigm can be applied in remote sensing, covert applications, and biological or biomedical imaging that utilize damaging or quantum light.

## Introduction

Despite considerable advances in multi-omic methods, optical microscopy remains the gold standard for non-invasively and spatiotemporally probing the physiology of living organisms.^1–5^ However, standard microscopy techniques come with a key challenge: the energy of the optical excitation field is only weakly transferred to the specimen during imaging.^6, 7^ Inevitably, this results in the need for higher irradiance levels, which during hour-long 4D imaging can considerably exceed the amount of sunlight to which life on Earth has adapted.^8^ While it remains challenging to quantify irradiance effects^9^, it suffices to say that irradiance levels higher than the solar flux has the potential to impair specimen physiology, or even kill it, leading to considerable changes in the conclusions drawn from an experiment.^6, 7^

To date, several approaches have been deployed to reduce specimen irradiance in optical imaging.^6^ One is light-sheet microscopy^10–15^ that elegantly confines the irradiance within the imaging plane, thereby minimizing unnecessary energy deposition on the specimen. Another approach involves sensitive photodetectors.^16–19^ In this context, high-quality scientific cameras with no amplification require approximately 10^3^ signal photons per pixel per frame to overcome the shot-noise limit and provide sufficient contrast (**Fig. 1a**).^20^ In contrast, sensitive detectors operating at the shot-noise limit, such as photon-counting or select electron-multiplying devices^16–19^, require approximately 10^2^ signal photons per pixel per frame (**Fig. 1b**). These signal fluxes may seem insignificant; however, they correspond to higher irradiance levels than one solar constant^8^ and, as such, can be toxic, especially for species that have not adapted to direct sunlight (e.g., rhizosphere microbiomes^21^).

**Figure 1.**
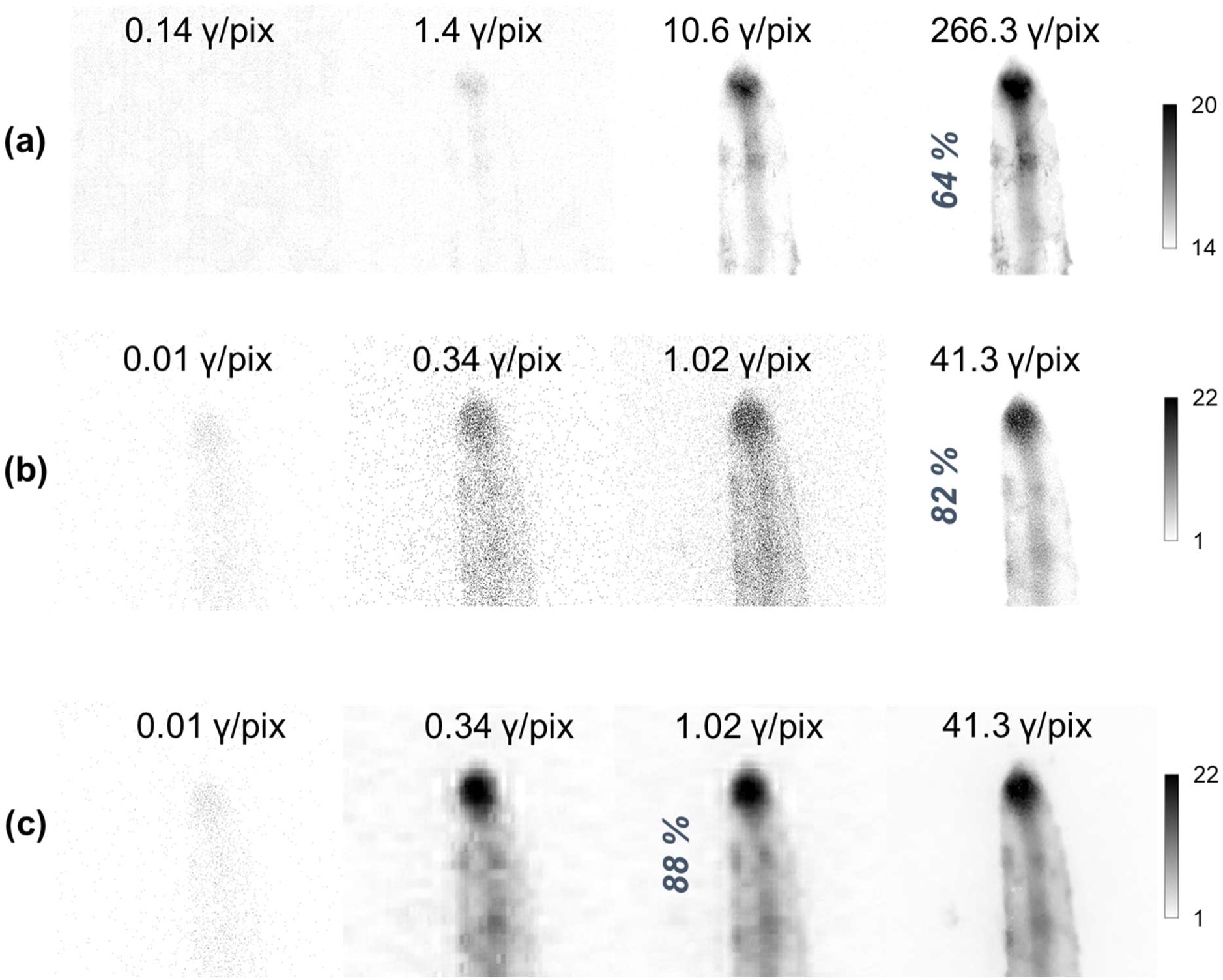
Low-light imaging. Epi-fluorescence images of a *M. truncatula* root at increasing photon fluxes using conventional **(a)** and photon-counting **(b)** detectors. A 64% contrast at 266.3 photons (γ) per pixel can be achieved using a conventional detector **(a)**, while photon-counting achieves 82% contrast at 41.3 photons per pixel **(b)**. Denoising by discrete wavelet transforms (**Methods**) of photon-counting images enables higher contrast (88%) at lower photon fluxes (1.02 photons per pixel) **(c)**.

Clearly, identifying ways to further reduce the energy deposited on a specimen during 4D imaging below one solar constant continues to be an important biotechnology goal. Further irradiance reductions, however, are met with the challenge of Poisson noise, or photon-sparsity. In this context, as irradiance is reduced, the particle nature of light is heightened, which increases detection uncertainty and enforces most image pixels (> 99%) to remain dark (**Fig. 1b**). Photon sparsity can be addressed by extending the acquisition duration, at the cost, however, of suboptimal imaging rates. Photon sparsity can also be addressed computationally; however, related algorithms^22, 23^ tend to fail at signal fluxes lower than 1 photon per pixel per frame (**Fig. 1c**), or – alternatively – at irradiance levels that are lower than one solar constant.

Here, we introduce *near-zero-photon imaging*, a method that operates at 1000× lower irradiance levels than one solar constant. Our approach combines a fluorescent microscope that acquires ultra-photon-sparse images at kHz rates with a supervised artificial intelligence approach that learns to eradicate Poisson noise and photon sparsity. Our method is different from others followed so far^23, 24^ in that it operates with classical light at the ultra-photon-sparse limit at near-zero-illumination conditions. As such, to the best of our knowledge, we are addressing for the first time the challenge of reconstructing classical ultra-photon-sparse images and validating our findings by imaging a wide variety of biological targets acquired under real experimental conditions.

We report that near-zero-photon imaging faithfully reconstructs the dark-pixels of photon-sparse images of both intensity and morphological biological features from as low as 0.01 photons per pixel. These near-zero-illumination conditions correspond to 10,000× reduction in irradiance levels compared to modern optical microscopy. These significant irradiance reductions are achieved at kHz imaging rates, thus violating the longstanding microscopy trade-off between toxic illumination and slow imaging, an important step towards practical, reliable, and reproducible bioimaging. As such, we anticipate that our method will accelerate biological and biomedical imaging, but also remote sensing, covert, and quantum applications.

## Results

To develop and validate near-zero-photon imaging, we deployed a judiciously designed laser-based epifluorescence microscope. As displayed in **Fig. 2a** and further detailed in the **Methods** section, the microscope was equipped with a CMOS camera that operated at 3.8 kHz. Further, the CMOS camera was integrated with an intensifier (iCMOS) to enable single photon detection at the Poisson limit (**Methods**). As per previous photon-sparse demonstrations^25, 26^, such semiclassical operation is largely independent of specimen and magnification, and we have observed both in fluorescence and Raman imaging. As such, we anticipate our findings described below to be valid not only for fluorescence imaging, but also Raman and other microscopy modalities. It is also worth noting that the observed Poisson dynamics emanate primarily from the selected temporal acquisition conditions and the multimodal nature of the imaging process, rather than inferring coherent photonic states.^27^ Further, near-zero-photon imaging operates strictly with photon-sparse images, namely images with considerably low background levels, where sparse photons occupying a single pixel represent the signal. As such, any imaging sensor that can deliver such images is in principle compatible with near-zero-photon imaging, such as single-photon avalanche photodiodes (SPAD) or avalanche photodiodes (APD) arrays and select electron-multiplying CCD (EMCCD) cameras.^16–19^

**Figure 2.**
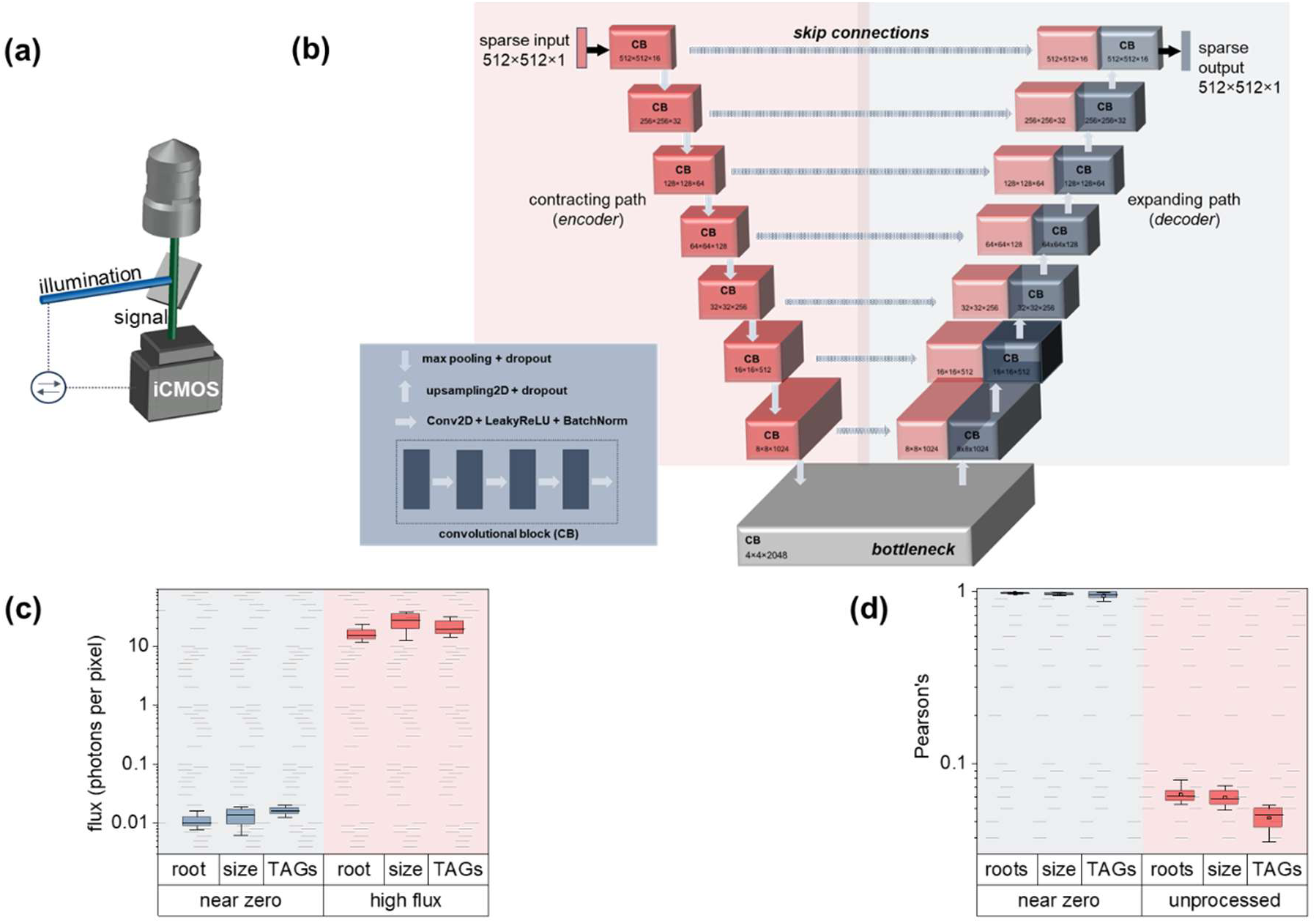
Near-zero-photon imaging experiment and learning model. The near-zero-photon epifluorescence setup **(a)** and U-Net architecture for image reconstruction, including the encoding and decoding paths linked with *skip connections* **(b)**; each path consists of 8 spatial tiers comprised of convolutional blocks (CBs) of four repeating convolutional layers; each layer uses a LeakyReLU(α=0.1) activation function followed by batch normalization; after each encoder CB, MaxPooling(2,2) downsampling is followed by a Drop-Out(0.5) layer to mitigate overfitting. **(c)** Photons per pixel for the near-zero and high-flux images for all biological samples and **(d)** the resulting Pearson’s coefficients between processed and unprocessed (raw) near-zero photon images with the ground truth; boxcharts and error bars represent 25% - 75%, and 10% - 90% ranges, respectively.

To suppress background-light, we modulated the illumination at 100 kHz by electrooptic means using the gating electronics of the intensifier as a trigger. Under these conditions, the intensifier fired ∼25 times per individual CMOS frame. Further, we cooled the intensifier down to -20°C, enabling ultralow dark photon counts per pixel per frame counts (6.5·10^−7^ ± 8.4·10^−8^, mean ± SEM for 7,000 CMOS frames). The electrooptic synchronization and ultralow dark count rates were key towards high contrast and error free image acquisition and reconstruction under near zero-photon conditions without adopting quantum light.^23^ Overall, this hardware configuration enabled the acquisition of super- and subcellular imaging data at magnification levels ranging from 4× to 100× (**Methods**).

### Learning model

A key of our approach is a deep learning system that reconstructs near-zero-photon images with exceptional efficiency and quality, preserving both brightness and spatial information. Our system is based on an optimally parameterized 2D convolutional autoencoder (CAE) and a specialized image processing workflow. Generally, CAEs learn to extract high-order features from data by first producing an encoded representation of the input image (encoding path) and then reconstructing the original image (decoding), while minimizing differences to the original input. Through this iterative process, CAEs can capture the most important spatial features of an image. U-Net architectures^28, 29^ are a form of CAE that are capable of retaining very fine spatial detail, thus finding tremendous success in biomedical image segmentation^30^ and denoising^31^, as well as generative text-to-image workflows^32^.

We found that conventional U-Net implementations yielded unsatisfactory reconstructions of near-zero-photon images; however, an exhaustive grid search of both hyperparameter space and network size revealed some key innovations that improved model accuracy. Specifically, we found that a U-NET architecture that was both *deeper* and *wider* than conventional implementations (**Fig. 2b**) greatly improved performance. Our implementation was *deeper* in that it consisted of eight spatial tiers, with each tier comprising a convolutional block built as a chain of convolutional layers. All layers within the first convolutional block used a filter size of 16, with each successive tier in the encoding path doubling the number of filters of the previous tier. In this way, the feature map at the U-Net bottleneck was relatively *deep* at 2048 layers. Our U-NET architecture was *wider* in that each convolutional block on every tier consisted of at least 4 successive convolutional layers. In this way, the network reached 250,384,297 parameters, increasing its ability to extract complex features. As per previous demonstrations, each successive tier in the encoding path used a max pooling operation^33, 34^ that doubled the size of the feature map while halving the spatial width and height of the tensor.

As input, our network takes 512×512 32-bit floating point tensors representing single photon images (**Methods**). This input was iteratively encoded through the convolutional blocks and max pooling layers to a bottleneck of 4×4 with a feature map that was 2048 layers deep. For decoding, we successively spatially upsampled using convolutional blocks of the same spatial dimensions per tier as the encoder, resulting in an output tensor with the same 512×512 dimensions as the original input. After each 2D upsampling step, we introduced skip connections^35, 36^ by concatenating the upsampled feature map with the feature map produced by the same spatial tier in the encoder. These skip connections represent a key innovation of our U-Net architecture, as they produced a feature map during decoding that combined the explicit spatial information from the encoder (prior to the bottleneck) with deeper features (extracted through the bottleneck). Further *widening* beyond four convolutional layers led to the deterioration of the model accuracy, while training and inference decelerated considerably, at rates that were approximately inversely proportional to the number of convolutional blocks.

Beyond *widening* and *deepening* the U-Net, we also made four key adjustments to mitigate model overfitting (**Methods**). In this context, we *first* performed batch normalization^37^ (**Fig. 2b**) after every convolutional layer to re-center and re-scale layer inputs. *Second*, we accompanied every max pooling or upsampling step with a dropout layer, which is known to mitigate model overfitting in convolutional neural networks.^38^ *Third*, we increased the size of the validation set to ensure that a larger and more diverse dataset evaluated model performance. *Finally*, we modified our approach to model selection by implementing a form of early stopping during training.^39^ This enabled us to adopt the best model based on validation performance instead of relying on the last model obtained during training, irrespective of epoch number. By retaining and using only the best model as measured through extensive validation during training, we greatly minimized any overfitting issues and improved the model’s overall generalization.

In addition to the 250,331,337 fully trainable parameters, our U-Net implementation consisted of 52,960 non-trainable ones, which played a key role in facilitating the functioning of the architecture. Importantly, our overall supervised training workflow used approximated ground truth images acquired through near-zero-photon images over longer exposure periods. This differs from conventional AI image reconstruction techniques that may require additional imaging modalities or synthetic data to establish ground truth examples; in contrast, our approach streamlined the training process, as well as reduced dependencies on additional image acquisition hardware and data resources.

### Near-zero validation

To validate the performance of near-zero-photon imaging, we collected and analyzed a diverse set of experimental bioimaging data. This dataset included targets ranging from multicellular organisms (*Medicago truncatula* roots, **Methods**) at lower magnification to cellular and subcellular features of single microbial cells at high magnification (*Yarrowia lipolytica* cell size and triacylglyceride – TAG – content, **Methods**). Given the inherently stochastic nature of CAE training, we evaluated a total of 720 near-zero-photon images to ensure the statistical significance of our findings. We collected images for all targets in two modes simultaneously: a *near-zero* and a *high-flux* mode. Near-zero images exhibited 0.01 ± 1.52·10^−4^ (mean ± SEM, **Fig. 2c** and **Fig. 3**) photons per pixel per frame and were deployed as tensor inputs to the CAE model. In contrast, high-flux images of the same targets were collected at ∼2,000× higher fluxes (20.96 ± 0.29 photons per pixel, **Fig. 2c**). This image dataset was used for training and validating the model results (**Methods**).

**Figure 3.**
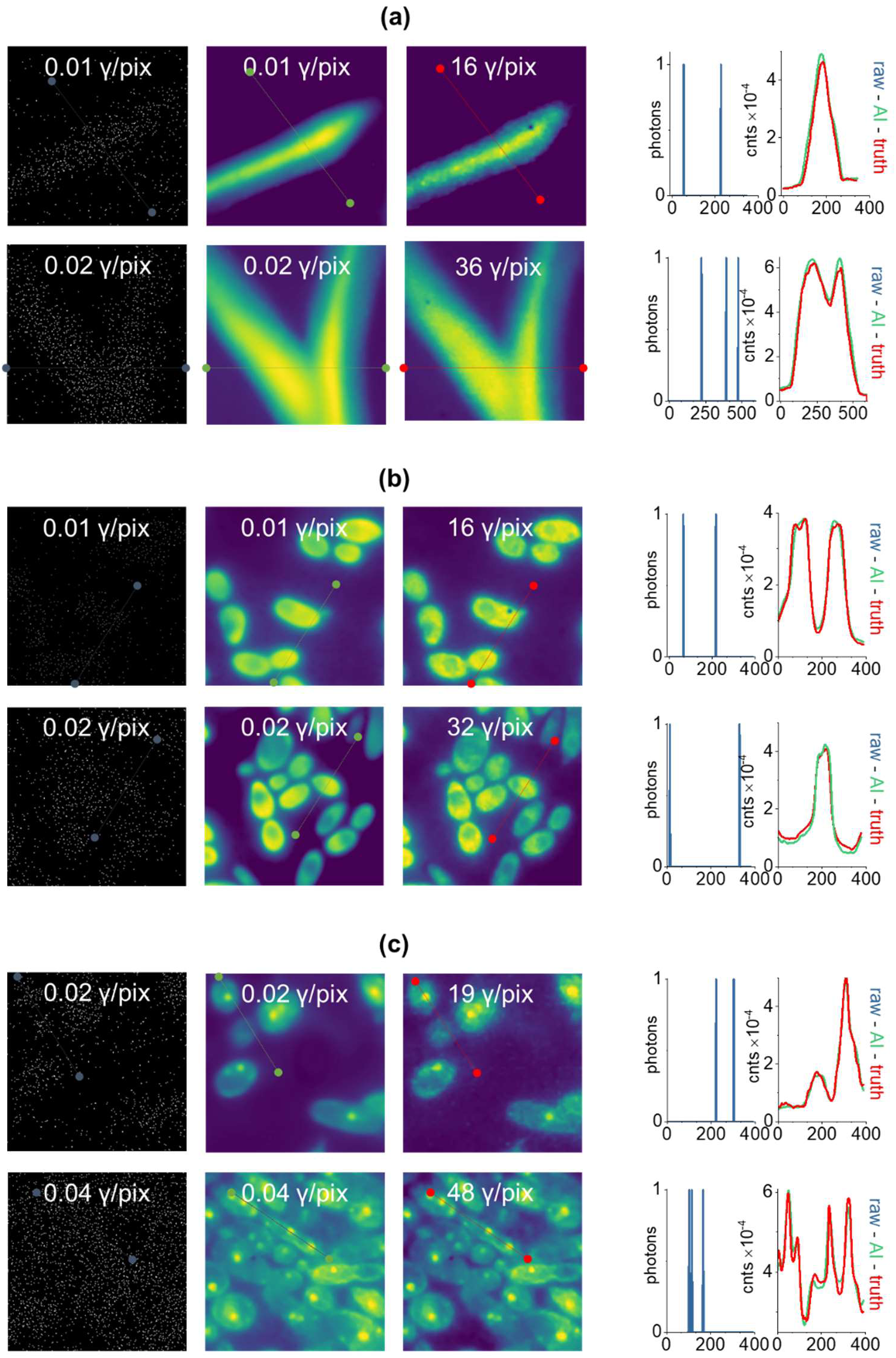
Near-zero-photon imaging of fluorescently stained *M. truncatula* roots **(a)**, *Yarrowia lipolytica* cells highlighting their size **(b)** and lipid (TAG) content **(c)**. All three cases display the raw (unprocessed, *1^st^ column*) and AI-reconstructed (*2^nd^ column*) near-zero-photon images, along with the respective high-flux ground-truth images (*3^rd^ column*). The *4^th^ column* compares exemplary 1D features in the reconstructed near-zero-photon and ground truth images that are represented by nearly zero photons in the raw image (each trace is noted in the respective images with dotted lines in *μm* units).

To quantitatively evaluate the performance of our proposed bioimaging method, we first assessed the similarity between unprocessed and processed near-zero images and their reconstructed, high-flux, counterparts (**Methods**). For these comparisons, we adopted the Pearson’s correlation coefficient metric, as per similar reports.^24^ Overall, the raw (non-reconstructed) near-zero images exhibited severe deterioration with almost no information content, as reflected by their low Pearson’s coefficient (0.06 ± 4·10^−4^, mean ± SEM, **Fig. 2d**) and Fourier spectra.^40^ In contrast, the morphological and intensity representation of the near-zero images after AI reconstruction were vastly improved, yielding a 16-fold higher Pearson’s coefficient (0.963 ± 0.001, **Fig. 2c**). This level of improvement is also apparent in the selected examples of **Fig. 3** and in alternative image similarity metrics, such as the structural similarity index. Additional examples of this performance are showcased in, including the occasional instances where the learning model introduced artifacts in the reconstructed near-zero-photon images. We also evaluated the uncertainty of the CAE model^40^, which exhibited a low coefficient of variation (CoV) of 6% or less across all tested examples.

It is important to highlight that we observed no resolution deterioration in the reconstructed near-zero photon images in comparison to their high-flux counterparts. To specifically demonstrate this, we trained our model to reconstruct images of dilute fluorescent particles, each 100 nm in diameter (**Methods**), imaged at a 100× magnification. The results demonstrate that the spatial resolution in the near-zero photon images is comparable to that in the high-flux images. This consistency across a range of photon flux levels confirms that the AI reconstruction process does not compromise the spatial resolution of near-zero photon imaging, thereby validating the efficacy of AI approach in maintaining image quality even under challenging ultralow-light conditions.

Finally, it is worth mentioning that in our investigation, low magnification root imaging yielded the highest average Pearson’s coefficient (0.978 ± 0.002, mean ± SEM), while intracellular lipid droplet imaging recorded the lowest (0.945 ± 0.004). Despite this, all imaging examples yielded Pearson’s coefficients considerably higher than those reported in similar reports^24^, while enabling us to conduct accurate morphological phenotyping across all magnifications, as further elaborated in the subsequent section below.

### Near-zero-photon phenotyping

To further verify the reliability and effectiveness of our method, we applied near-zero-photon imaging in phenotyping the same biological targets, ranging from multicellular organisms to cellular and subcellular features. In this context, we first imaged *M. truncatula* roots until a 0.15 μm^-1^ spatial frequency and a 0.0036 ± 3·10^−4^ average photon flux per pixel per frame. Through this analysis, we found that primary and secondary roots, and their vascular bundle can be effectively restored using our CAE model, even if they were previously indiscernible in the unprocessed near-zero-photon images. As per **Fig. 3a**, the fluorescence cross-sectional traces across the root tissue can be accurately reconstructed in excellent agreement with the high-flux images in terms of both intensity and root diameter (blue traces in **Fig. 3a**). Importantly, this pertains to features that are literally represented by *nearly zero* photons in the raw images. Higher throughput screening (41 different roots) similarly revealed excellent agreement in phenotyping root size (5.3 ± 0.4% error, mean ± SEM) with the *high-flux* images (**Fig. 4a** and **4b**).

**Figure 4.**
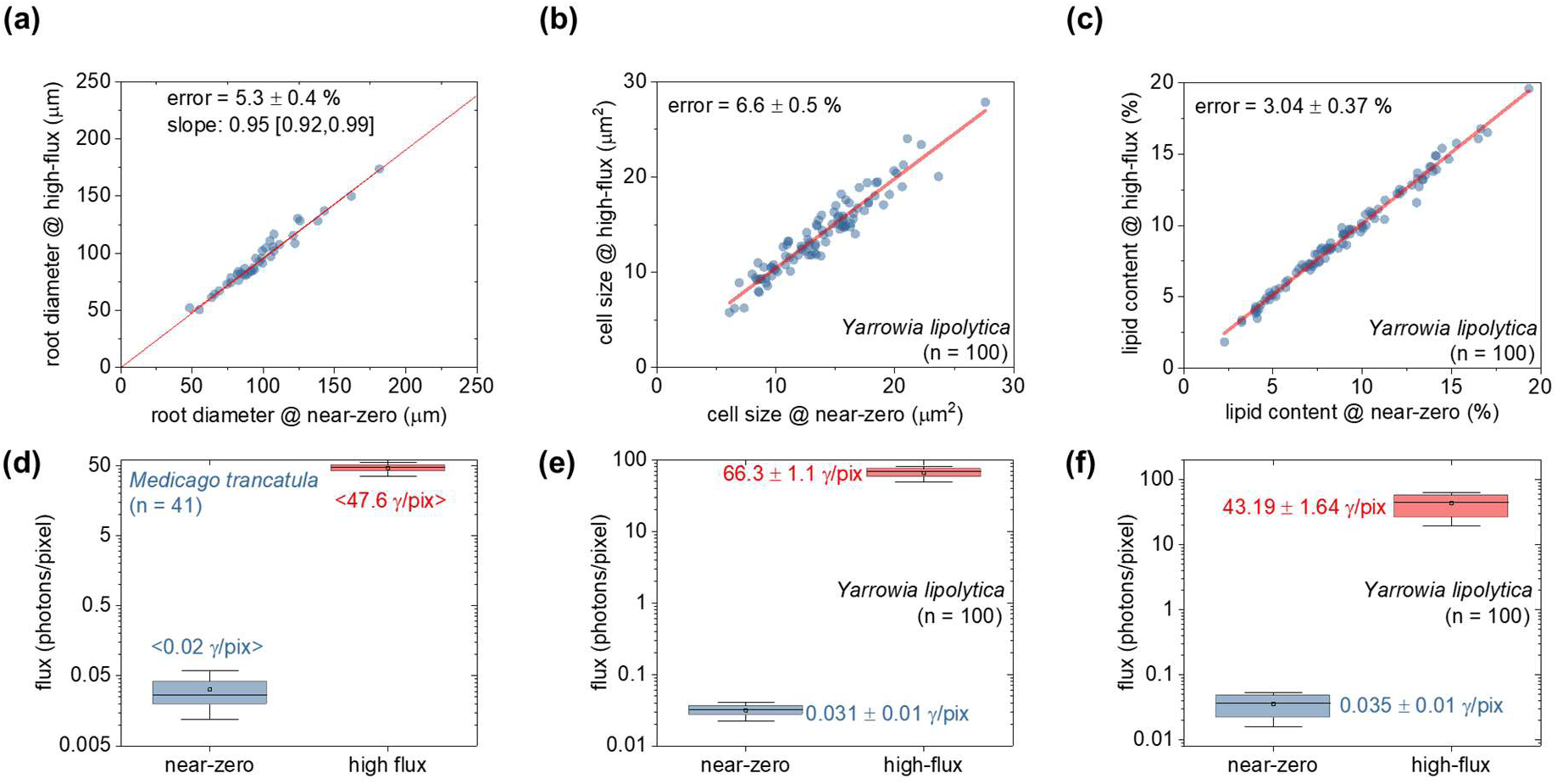
Near-zero phenotyping. Scatter plots comparing various phenotypic characteristics that were independently determined by near-zero (*x-axis*) and high-flux (*y-axis*) imaging. The phenotypic characteristics include: the diameter **(a)** of primary *M. truncatula* roots, *Y. lipolytica* cell size **(b)**, and *Y. lipolytica*’s lipid droplet (LD) % content **(c)**. Legends in all scatter plots detail the normalized (%) error between near-zero and high-flux conditions (mean ± SEM). The photon fluxes **(c-f)** for the near-zero and high-flux regions of interest (ROIs) used in phenotyping the root size **(a)**, the size **(b)** and the LD-content of *Y. lipolytica* cells **(c)**; boxcharts and error bars represent 25% - 75%, and 10% - 90% ranges, respectively.

We also validated near-zero-photon imaging in phenotyping at considerably higher spatial frequencies of up to 4.7 μm^-1^. In this context, we imaged the yeast *Yarrowia lipolytica* to find that both cell size and the intracellular TAG content can be successfully reconstructed using our CAE model from an average photon flux of 0.031 ± 0.01 and 0.035 ± 0.01, respectively. As shown in **Fig. 3a** and **3c**, one-dimensional cytosolic traces can also be restored in excellent agreement with the high-flux images in both intensity and morphology, even if they are represented by nearly-zero photons in the raw images. Higher throughput screening (100 single-cell observations) exhibited 6.6 ± 0.5 % and 3.04 ± 0.37% (mean ± SEM, **Fig. 4b,c**) in quantifying the cell size and TAG content. The analysis at increased spatial frequencies revealed that near-zero-photon imaging can be generalized with exceptional scalability and comparable performance, regardless of magnification levels.

### Specialization vs. Generalization

Near-zero-photon imaging, at its core, leverages deep learning, where accuracy and generalizability are strongly dependent on the quality, quantity, and variability of the data used for training and evaluation.^40^ The intrinsic low information content (entropy) of ultra-sparse photon data necessitates training of specialized models for each specific sample type or imaging condition to ensure optimal results. This focused training requirement is similar to other deep learning demonstrations in microscopy, such as recent low-light imaging^24^ and semantic segmentation^41^ applications. Overall, specialized datasets can hinder flexibility in some applications by requiring customized training to suit different imaging hardware configurations or sample types. Despite the need for specialized training, near-zero-photon imaging can be effectively used to capture live cell images that reveal time-dependent information.

Further, it is possible for some applications to pretrain models on larger datasets and adapt/transfer them to new tasks. We explored this possibility for near-zero-photon imaging by pretraining a single model on yeast cells stained with two different fluorophores: Bodipy (for visualizing intracellular TAGs) or Propidium Iodide (for visualizing cell size). The model trained on cells stained with both fluorophores achieved an average Pearson’s coefficient of 0.97 ± 0.01 (mean ± SEM), which is equal to models trained exclusively on cells stained with either one of the fluorophores (0.97 ± 0.01). These findings indicate that while more targeted training achieves higher accuracy, it is possible to generalize nearzero-photon imaging towards reconstructing images across more than one imaging condition and/or target type with strong results. We observed a similar effect in near-zero photon images of live *Saccharomyces cerevisiae* cells expressing a red fluorescent protein (RFP) were successfully reconstructed using a model pre-trained in fixed *Y. lipolytica* cells stained with propidium iodide.

## Discussion

The broad spectrum of experimental samples and questions targeted by biological or biomedical imaging challenges the precise quantification of the deleterious effects of standard irradiance levels^9^ or the development of standardized assays to quantify them.^6^ It suffices to say, however, that standard microscopy adopts irradiance levels that exceed the level of natural sunlight to which life on Earth has adapted.^6^ This concept becomes apparent upon considering the efficiency at which optical irradiance is converted to practical imaging information. Using 1-photon epifluorescence imaging of gene-edited chromophores^42^, a common contrast mechanism in biological imaging, as an example, we deduce that this excitation efficiency depends on:

- the quantum yield of the fluorophore (∼60 % on average^43^);
- the weak absorbance in a cell (∼6.03·10^−5^ for a yeast cell expressing an exorbitant amount of 100,000 protein copies^43^);
- the imperfect collection efficiency of commercial optical components (50% max, assuming a point source with isotropic emission);
- the constrained quantum yield of most photodetectors (∼ 90%).

Assuming an imaging experiment using advanced photodetectors operating at 10^2^ signal photons and 20 Hz rates, we deduce that these experiments would require approximate illumination levels of 10^7^ photons per pixel per frame. These fluxes are 10× higher than one solar constant^8^ and, as such, can be toxic to most organisms. Further, these fluxes increase by more than a factor of 10^3^, when executing hour-long imaging, as needed to satisfactorily probe the 4D cellular and subcellular dynamics. They also increase by ∼10^7^ per frame in alternative imaging contrast mechanisms, such as multiphoton or nonlinear microscopy^1^ and coherent or spontaneous Raman imaging^44^.

Near-zero-photon imaging, the method that we present here, can extract information without sacrificing image quality at 1000× lower irradiance levels than one solar constant, or approximately 10,000× lower irradiance than modern optical imaging methods. To attain this substantial advantage, our method deploys a judiciously designed epifluorescent microscope enabling practically zero background, and supervised learning of the innate spatial correlations that are present in any biological sample. Similar learning operations on nearly zero information inputs have been recently reported in a non-bioimaging context.^45, 46^ Our approach reduced irradiance at kHz imaging rates without being constrained by photon sparsity, which violates a longstanding microscopy trade-off between toxic illumination and slow imaging. Finally, it is worth mentioning that near-zero-photon imaging can confer key advantages against photobleaching, a key metric in time-lapse imaging. To illustrate this, we compared the fluorescence intensity dynamics of near-zero-photon imaging with conventional epifluorescence on fixed yeast *Y. lipolytica* cells. Notably, the excitation power in near-zero-photon imaging was 56,000× lower than conventional epifluorescence, at 1 kHz rates (limited by the sCMOS frame-rate). Under these conditions, near-zero-photon imaging exhibited virtually no photobleaching, while conventional epifluorescence lost 50% intensity in 6 minutes. Finally, near-zero photon imaging is congruent with but does not require quantum correlations that can be limited by optical losses impeding the full realization of the quantum metrology benefits.^47, 48^

The reported irradiance reduction level of 10,000× compares well against recently reported artificial intelligence denoising methods conferring ∼10× irradiance reduction levels.^24, 49^ We attribute this considerable improvement to the photon-sparse operation of our microscope with essentially zero background noise. Importantly, our non-canonical CAE can be deployed for inference in state-of-the-art desktop machines and operate at 11 fps rates, which is close to real-time operation. Further, we demonstrated near-zero-photon imaging using standard epifluorescence as a contrast mechanism. As our approach relies strictly on the innate spatial correlations of any biological specimen, we anticipate full compatibility with alternative contrast mechanisms and greater irradiance reductions when near-zero-photon imaging is combined with advanced hardware, such as light-sheet illumination.^25, 26^ Overall, we expect that the paradigm of near-zero-photon-imaging will eradicate phototoxicity and photobleaching, an important advancement towards reliable biological and biomedical imaging. Near-zero-photon imaging can be improved by advances in single-photon detector technology.^18^

## Acknowledgments

We gratefully acknowledge funding from the U.S. Department of Energy, Office of Science, Office of Biological and Environmental Research (DE-SC0022282). We also acknowledge fruitful discussions on epifluorescence imaging with Dr. Ramachandran Kasu.

## Author contributions

LS: development, validation, and execution of deep learning models; SB: experimental data acquisition; MJH: provided *M. truncatula* materials, growth methods and image interpretation (root structure); JJ provided the *S. cerevisiae* materials and image interpretation; AEV: design/overview of study, hardware development; manuscript writing with input from LS and MJH, funding acquisition.

## Data availability statement

The data and software that support the findings of this study are available online.

